# Insight into the autosomal-dominant inheritance pattern of SOD1-associated ALS from native mass spectrometry

**DOI:** 10.1101/2020.07.13.199976

**Authors:** Jelena Cveticanin, Tridib Mondal, Elizabeth M. Meiering, Michal Sharon, Amnon Horovitz

## Abstract

About 20% of all familial amyotrophic lateral sclerosis (ALS) cases are associated with mutations in superoxide dismutase (SOD1), a homodimeric protein. The disease has an autosomal-dominant inheritance pattern. It is, therefore, important to determine whether wild-type and mutant SOD1 subunits self-associate randomly or preferentially. A measure for the extent of bias in subunit association is the coupling constant determined in a double-mutant cycle type analysis. Here, cell lysates containing co-expressed wild-type and mutant SOD1 subunits were analyzed by native mass spectrometry to determine these coupling constants. Strikingly, we find a linear positive correlation between the coupling constant and the duration of the disease. Our results indicate that inter-subunit communication and a preference for heterodimerization greatly increase the disease severity.

---

Homodimeric proteins are widespread in Nature including in humans where they constitute about 25% of all proteins with a known crystal structure (1). Mutations in homodimers are often associated with disease (2). Examples include the involvement of Park7 (DJ1) in Parkinson’s disease (3), NAD(P)H quinone oxidoreductase 1 (NQO1) in cancer and certain neurological disorders (such as multiple sclerosis and Alzheimer’s disease) (4) and Cu, Zn-superoxide dismutase (SOD1) in amyotrophic lateral sclerosis (ALS) (5). Patients with the disease-causing mutations are often heterozygous. In other words, the mutation is found in only one of their two alleles and the disease follows an autosomal-dominant inheritance pattern, as in the large majority of SOD1-associated ALS cases (6). Consequently, patients may have three co-existing dimeric forms of the protein, which respectively contain either (i) two wild-type subunits (wt/wt); (ii) two mutant subunits (mut/mut); or (iii) one wild-type subunit and one mutant subunit (wt/mut). In principle, the wild-type and mutant subunits can associate randomly (i.e. with equal affinities) so that the wt/wt, wt/mut and mut/mut complexes are formed at a 1:2:1 binomial ratio, respectively, if the wild-type and mutant subunit concentrations are equal (i.e. the probabilities for formation of the wt/wt, wt/mut and mut/mut complexes are ½ x ½, 2 x ½ x ½ and ½ x ½, respectively, which correspond to a 1:2:1 ratio). Alternatively, the wild-type and mutant subunits can associate preferentially so that the ratio of the wt/wt, wt/mut and mut/mut complexes deviates from the binomial expectation. In the work described here, we investigated whether the extent of deviation from random association of wild-type and mutant SOD1 subunits correlates with measures of clinical severity in ALS patients, such as disease duration after onset (7). Our approach should be applicable to many other diseases associated with mutations in homodimers (2).

The extent of deviation from random association of wild-type (wt) and mutant (mut) subunits can be quantified using a double-mutant cycle type of analysis (8). In its implementation here, these cycles consist of the wt/wt, mut/mut and the two equivalent wt/mut and mut/wt dimeric forms of the protein (Fig. 1). The change in the free energy of dimer formation upon a mutation in one subunit may be expressed relative to the free energy of formation of the wild-type dimer as Δ*G*(wt/wt→wt/mut). Likewise, the change in the free energy of dimer formation upon a mutation in one subunit, when the other subunit already has that mutation, may be expressed as Δ*G*(wt/mut→mut/mut). The difference between Δ*G*(wt/wt→wt/mut) and Δ*G*(mut/wt→mut/mut), which is equal to the difference between Δ*G*(wt/wt→mut/wt) and Δ*G*(wt/mut→mut/mut), provides a measure of the coupling energy between the mutated residues in the two subunits. This coupling can be expressed in terms of binding constants as follows:

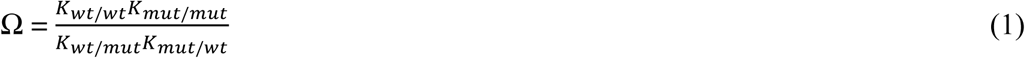

where *K*_wt/wt_, *K*_mut/mut_, *K*_wt/mut_ and *K*_mut/wt_ are the equilibrium constants for the formation of the four dimers (it should be noted that *K*_wt/mut_ and *K*_mut/wt_ are equal to each other). Given that each of the equilibrium constants in Eq. 1 is equal to the ratio between a dimer concentration and its corresponding subunit concentrations (e.g. *K*_wt/mut_ = [wt/mut]/[wt][mut]), the expression for the coupling constant simplifies to:

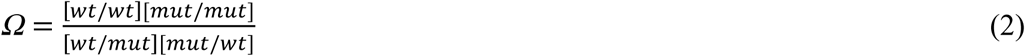

**Figure 1.**
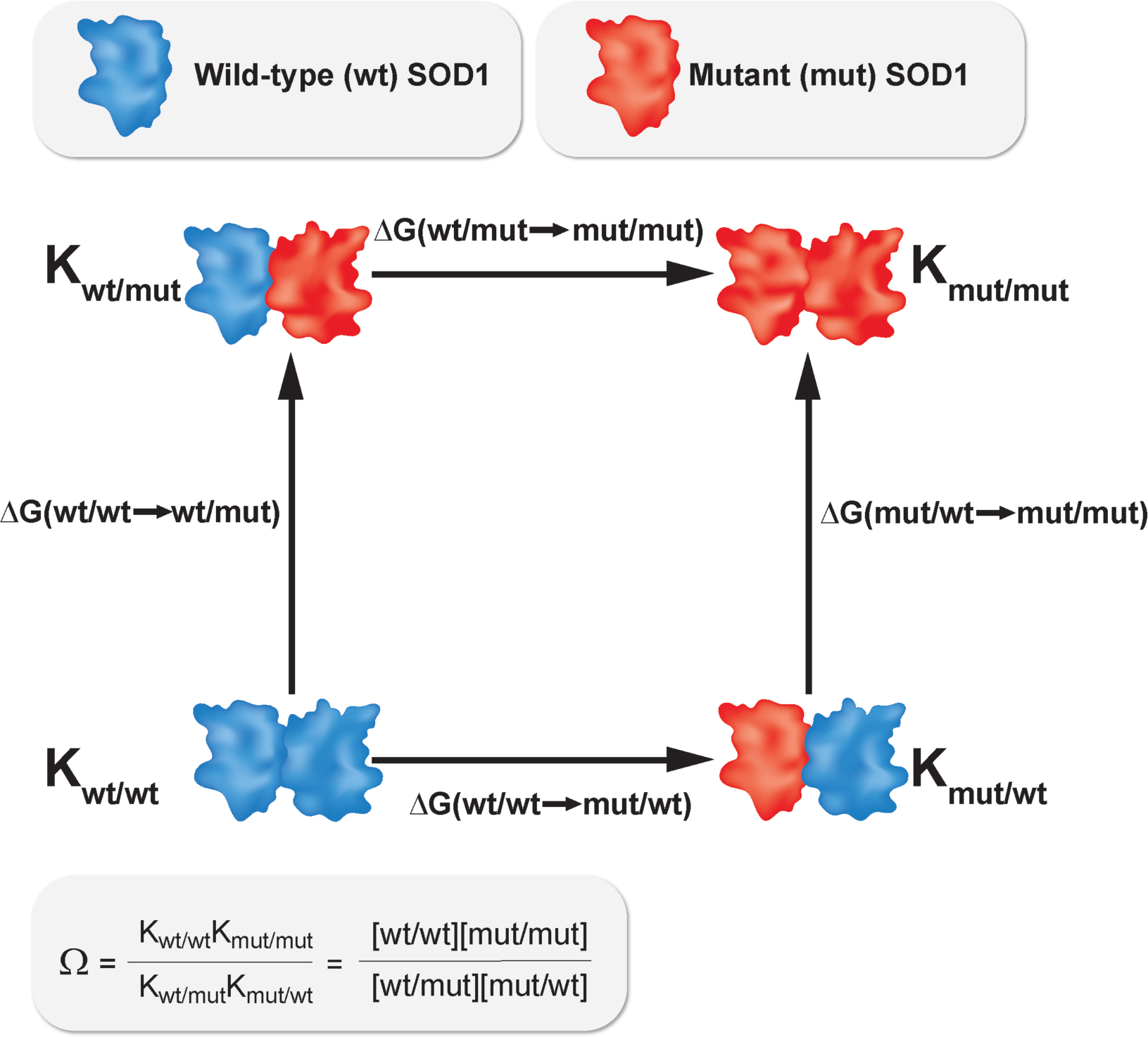
Double-mutant cycle for measuring coupling between wild-type and mutant SOD1 subunits. The cycle consists of the wild-type homodimer (wt/wt), the mutant homodimer (mut/mut) and the two equivalent heterodimers (wt/mut and mut/wt). The binding constants, *K*, corresponding to subunit association for the four dimers in the cycle and the changes upon mutation in the free energies of dimer formation are also indicated in the scheme. It should be noted that Δ*G*(wt/wt→wt/mut) = Δ*G*(wt/wt→mut/wt) because wt/mut and mut/wt are equivalent.

An Ω value of 1 indicates random association of wild-type and mutant subunits (i.e. self- and hetero-association have equal affinities) whereas Ω > 1 and Ω < 1 indicate preferential formation of both homodimers or the heterodimers, respectively. In other words, Ω provides a measure of the coupling energy between the wild-type and mutant subunits. Recently, we showed that one can estimate the values of such coupling constants by co-expressing the wild-type and mutant variants of two interacting proteins (or, as we now demonstrate here, the wild-type and mutant subunits, wt and mut, which form the homo- and heterodimers) and then determining the concentrations of all the co-existing complexes from a single native mass spectrometry (MS) spectrum (9). This is possible because native MS is based on the ability to transfer protein complexes to the gas phase without disrupting them (10-12). Moreover, such transfer can take place from crude cell lysates (13). This approach (Fig. 2), therefore, bypasses the requirement for protein purification, which can lead to perturbations in the structure and/or assembly of the proteins. It also obviates the need to determine the values of the individual equilibrium constants, which is often labor-intensive and error-prone. Importantly, coupling constants determined for crude cell lysates and purified proteins were found to be similar (13), thereby indicating that our approach does not perturb the equilibrium distribution *in vivo* of the various co-existing species.

**Figure 2.**
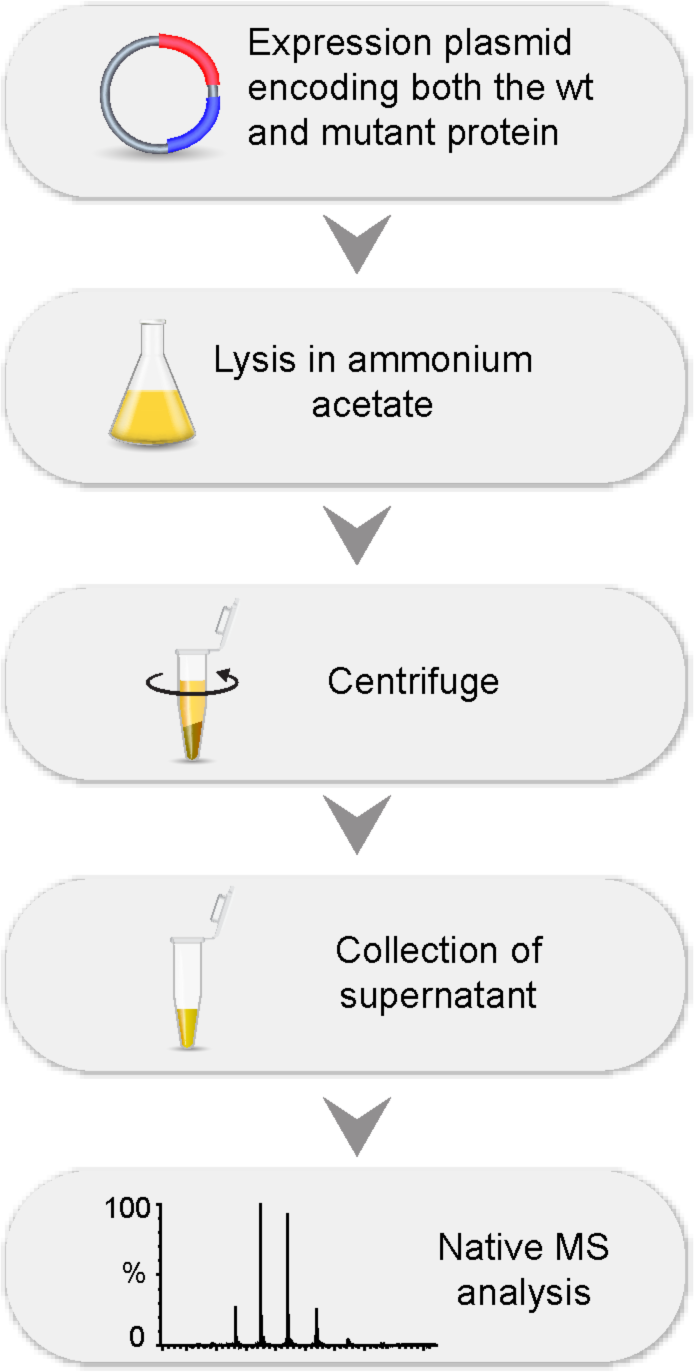
Flowchart depicting our experimental procedure. Wild-type and mutant SOD1 were co-expressed and the cells were then lysed in buffer compatible with native MS analysis. The lysates were then centrifuged and the supernatants were subjected to native MS analysis.

We chose to study ALS-causing mutations in SOD1 (i.e. A4V, G37R, E100G, H43R, G85R, G93S and L144F, Fig. 3) with diverse structural effects and which are associated with a wide range of average ages of onset and disease durations (14-16). The average disease duration, i.e. the time from disease onset to death, of patients with the mutation A4V, for example, is 1.2 years whereas that of patients with the mutation G37R is 17 years (7). These mutations are also relatively common so that estimates of their associated ages of onset and disease durations are most reliable. Consequently, many of these SOD1 mutants have been the focus of previous biophysical studies (17, 18). Wild-type SOD1 and each of the mutants, in turn, were co-expressed in *E. coli* so that both homodimers and the corresponding heterodimer could form at the same time. The cells were then lysed and the lysates were analyzed using native MS (Fig. 2). For all the mutants, it was possible to resolve the peaks corresponding to the wild-type and mutant homodimers and the corresponding heterodimer and determine their relative populations from their intensities (Figs. 4 & S1).

**Figure 3.**
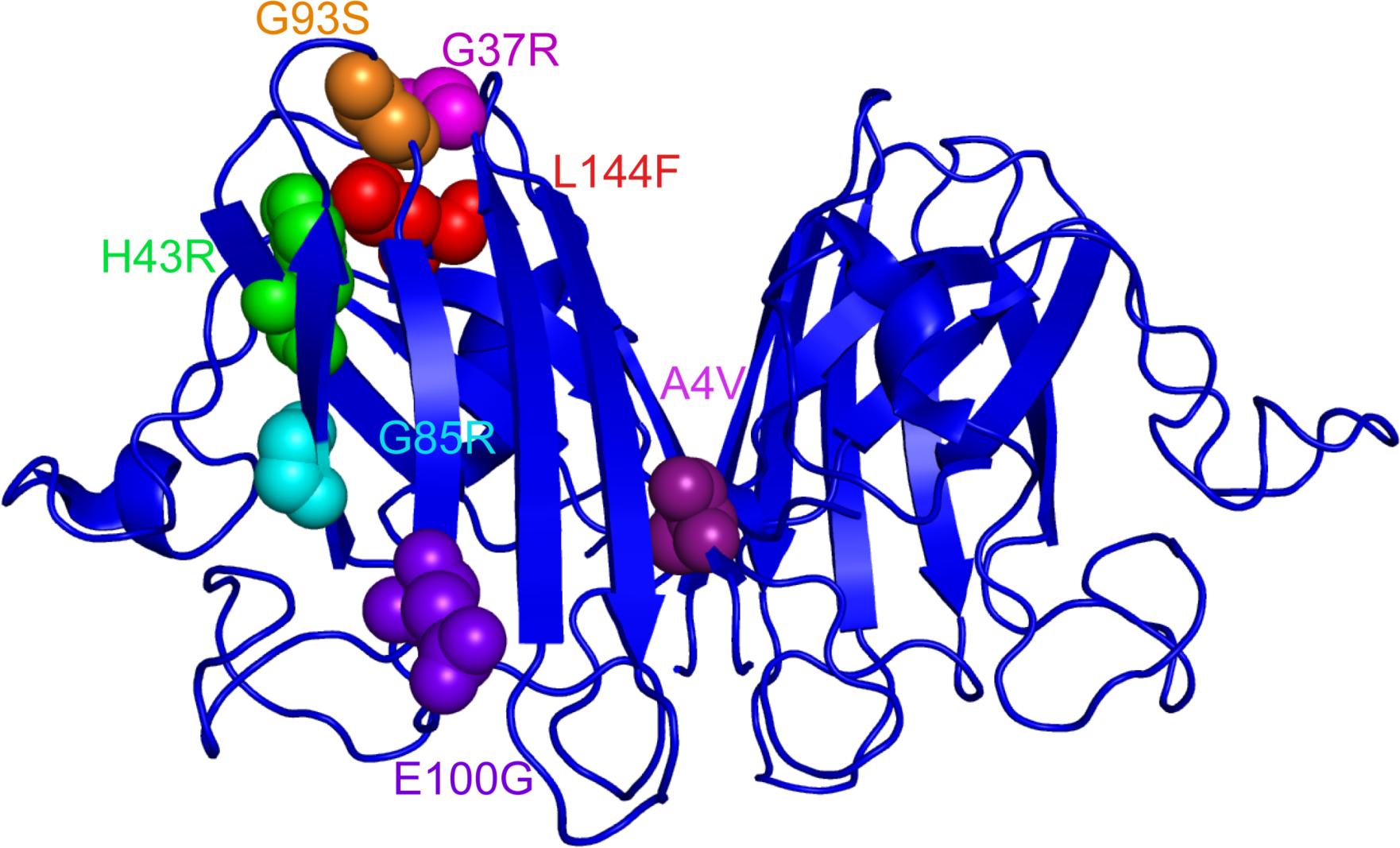
Ribbon diagram of the crystal structure of the pseudo wild-type SOD1 dimer containing the mutations C6A and C111S (PDB ID: 1n18). The mutations studied here are indicated in the left subunit next to the respective mutated residues using the single-letter code for amino acids. The figure was generated using PyMOL.

**Figure 4.**
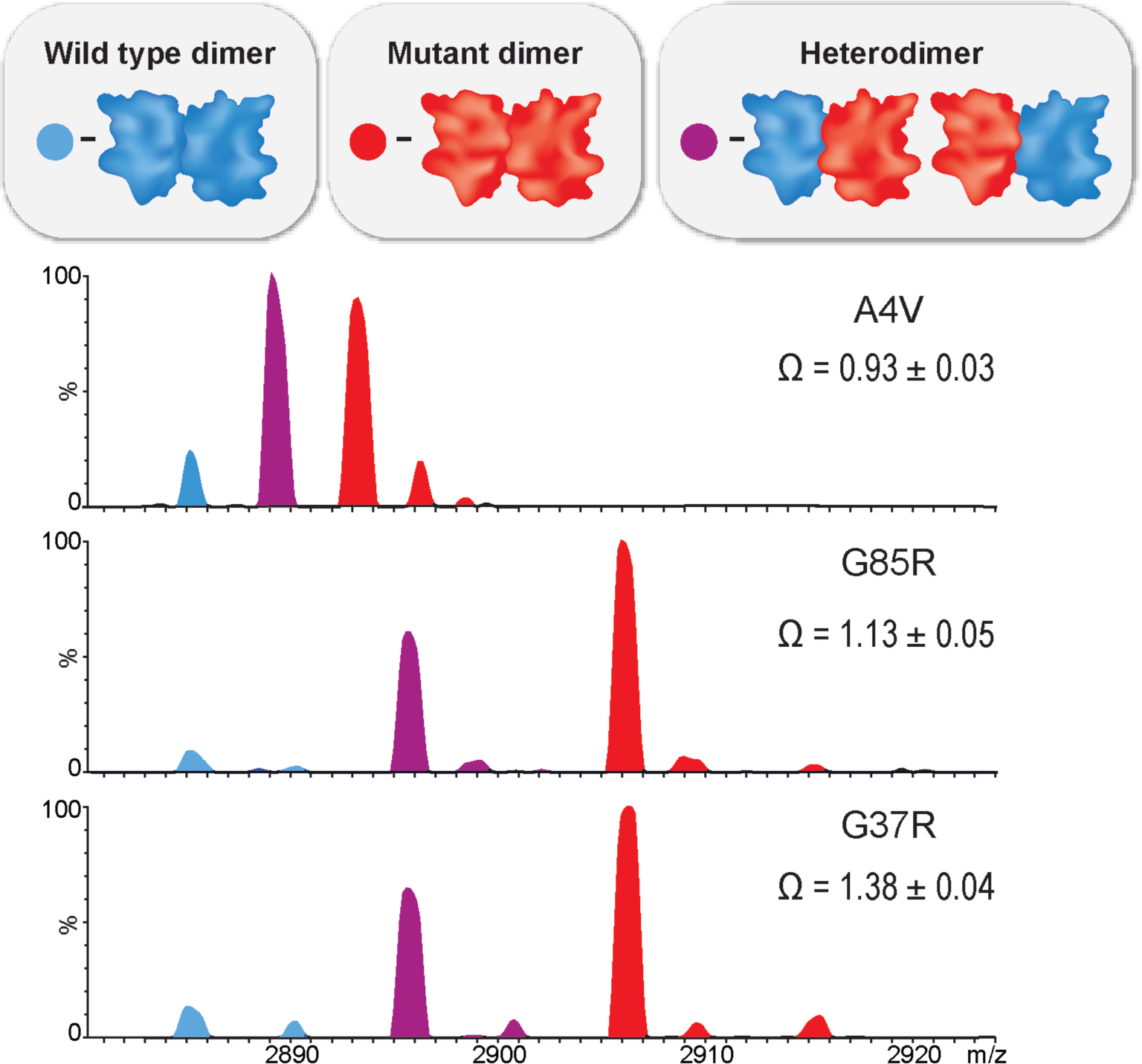
Native MS measurements of the wild-type and mutant SOD1 homo- and heterodimers in crude cell lysates. Representative mass spectra of the 11^+^ charge state are shown for lysates containing co-expressed wild-type SOD1 and either the A4V, G85R or G37R mutants. The peaks for the wt/wt, mut/mut and mut/wt (and wt/mut) dimers are shown in blue, red and purple, respectively. The adducts with masses of 36, 54 and 72 Da likely correspond to two, three and four NH_4_^+^ ions, respectively, and the adduct with a mass of 99 Da to a phosphate ion.

Strikingly, a strong positive linear correlation (r^2^ = 0.807; *P* = 0.006) is observed between the coupling constants of the different mutants and their associated disease durations (Fig. 5). Previously, a correlation was observed between the free energy of heterodimerization and disease duration for a different set of SOD1 mutants in their disulfide-intact apo form (17), which is little populated in ALS patients. Here, however, we assessed the metallated, mature form of SOD1, which is likely to be significantly populated in patients (19). Heterodimers have also been the focus of *in vivo* studies, which have highlighted their toxicity (20-23). Importantly, the analysis here using the coupling constant takes into account also the free energies of formation of the two homodimers (Fig. 1). This is important because mutations that affect *K*_wt/mut_ (heterodimer formation) are also likely to have effects on *K*_wt/wt_ and *K*_mut/mut_, which are taken into account in the calculation of the coupling constant (Eq. 1) as discussed further below. It is noteworthy in this regard that we also observe a correlation between the coupling constant and age of onset when the data for G37R are excluded (Fig. S2). By contrast, no correlation was observed in previous work between the free energy of heterodimerization and age of onset (17).

**Figure 5.**
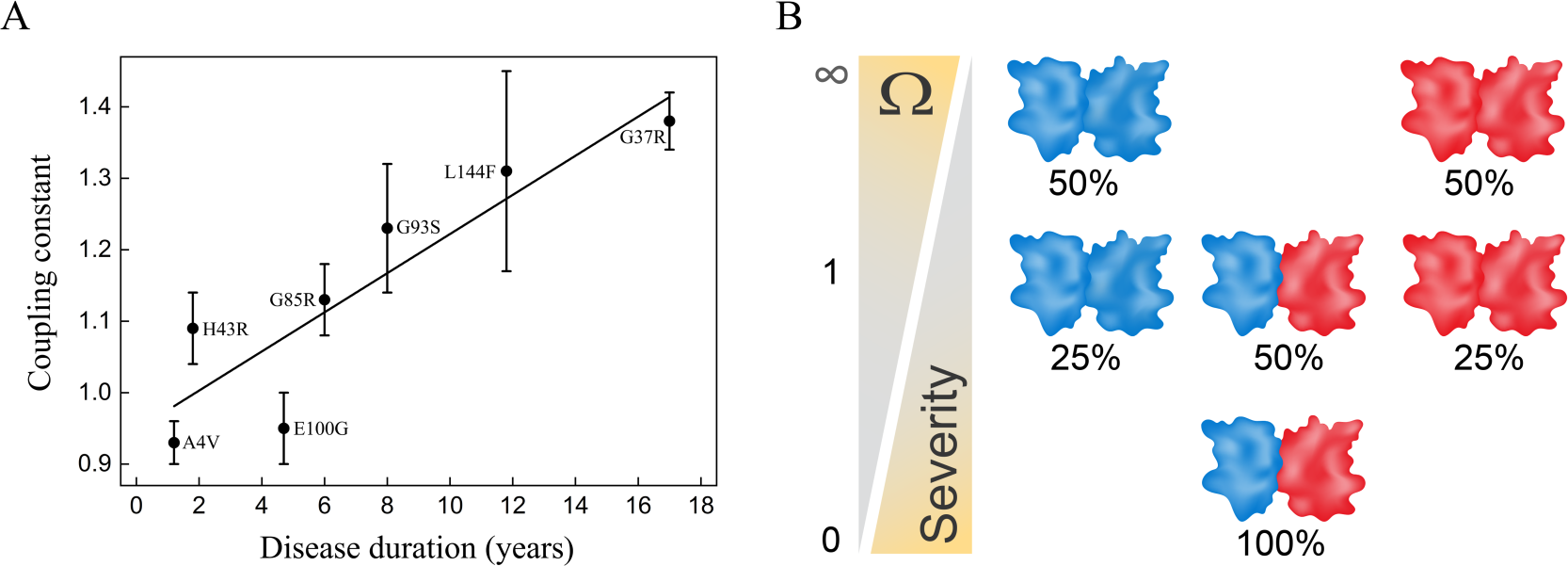
Disease duration correlates with coupling energy. (A) Plot of the coupling energies between wild-type and mutant SOD1 subunits and the disease duration associated with the respective mutation. For simplicity, the data were subjected to a linear fit (*r*^2^ = 0.807; P = 0.006) although the underlying relationship may be monotonically increasing but nonlinear. The disease durations were taken from ref. 7. (B) Scheme showing for the case of equal concentrations of wild-type and mutant subunits how the extent of preferential association is reflected in the value of the coupling constant.

The correlations we observe (Figs. 5 & S2) can be rationalized, in part, by considering how the fraction of dimers containing at least one mutant subunit depends on the value of the coupling constant, Ω. We illustrate this dependence for the case where wild-type and mutant subunits are present at equal concentrations. It is important to note, however, that the value of Ω is independent of the relative subunit concentrations (9). In the absence of coupling (Ω = 1), the wild-type and mutant subunits associate randomly and, therefore, 75% of the dimers (i.e. the 50% heterodimers and the 25% mutant homodimers) will contain at least one mutant subunit when the concentrations of both types of subunits are equal. In the case of infinite positive coupling (Ω = ∞), only homodimers form and, therefore, 50% of the dimers will contain at least one mutant subunit when the concentrations of both types of subunits are equal. Finally, in the case of infinite negative coupling (Ω = 0), only heterodimers form and, therefore, 100% of the dimers will contain at least one mutant subunit. In other words, the positive correlation we observe between the value of Ω and disease duration implies that the disease severity increases dramatically when the fraction of dimers containing at least one mutant subunit increases. The coupling constant is, however, superior to the fraction of dimers containing at least one mutant subunit as a measure of inter-subunit interactions because it is independent of differences in subunit concentrations (as mentioned above) and of the relative stabilities of the wild-type and mutant monomers (since their unfolding free energies cancel out in the double-mutant cycles here).

The coupling constants measured here provide insights into the effects of ALS-associated mutations in SOD1, which cannot be gained through other measurements. Many ALS-related mutations in SOD1 decrease its overall thermodynamic stability by reducing monomer stability (18, 24), interfering with metal binding (25, 26) and/or disfavoring dimerization (17, 27, 28). An extensive computational analysis of 75 ALS-linked SOD1 mutations showed that 70 of them increase dimer dissociation (28), thereby suggesting that this may be a major mechanism. Regardless of the mechanism causing destabilization, all these mutations can lead to misfolding, formation of small aggregates and ultimately the accumulation of insoluble inclusions, which are found in neurons of patients (29). Attributing the impact on disease severity of SOD1 mutations to their effects on stability ignores, however, the observed toxicity of mutant and wild-type heterodimers (20-23). It was suggested that such toxicity can be quantified by the free energy of heterodimer formation (17) but this measure neglects potential effects of the mutations on the stability of the co-existing mutant and wild-type homodimers. The coupling energies determined here take into account the stabilities of all the potentially co-existing dimers and are independent of monomer stability and expression levels. Therefore, they isolate the impact of inter-subunit coupling from other effects.

The correlation observed here between the coupling energy and disease duration (and to a lesser extent age of onset) highlights the tendency for heterodimerization and the role of inter-subunit communication in SOD1 in determining the disease severity. Such communication may lead to greater toxicity of wild-type subunits through their interaction with mutant subunits as proposed before (23) via a mechanism that will be of interest to explore. Our approach will be useful in the future for determining if such a tendency affects the severity of other diseases caused by mutations in homodimers (2) such as, in addition to the examples mentioned above (3-5), many transcription factors including the hepatic nuclear factors 1α and 4α in which mutations cause maturity-onset diabetes also with autosomal-dominant inheritance (30).

## Materials and Methods

### Mutagenesis and cloning

The disease-causing mutations H43R, G85R, G93S and L144F were introduced into the gene coding for pseudo-wild-type SOD1 in a pBR322-derived plasmid using the restriction-free (RF) cloning method (31). The pseudo-wild-type SOD1 gene contains the replacements C6A and C111A (32) and is referred to throughout this paper as wild-type SOD1. Construction of the mutations A4V, G37R and E100G has been described before (33). RF cloning was also used to introduce the pseudo-wild-type SOD1 gene into the pRFSDuet-1 plasmid (Novagen). The mutant SOD1 genes were amplified with primers containing restriction sites for NdeI and AgeI and then inserted into the pRFSDuet-1 plasmid using the In-Fusion® HD cloning kit after digesting the plasmid with the same enzymes. All the mutants also contain the mutation C111S instead of C111A in order to increase the mass difference between them and the wild-type protein. The disease-causing mutations H43R and G93S were also accompanied by addition of a C-terminal alanine in order to further increase the mass difference between these mutants and the wild-type protein. Control experiments showed that addition of a C-terminal alanine has a relatively minor effect on the coupling constant of the G37R variant (1.38 ± 0.04 vs. 1.28 ± 0.01). The mutations were confirmed by DNA sequencing of the entire genes and by the mass spectrometry results. The following forward mutagenic oligonucleotides were used (an asterisk follows the mismatched bases):

H43R: 5’-CATTAAAGGACTGACTGAAGGCCTGAGAGGATTCCATGTTCATGAGTTTGGAGA-3’

G85R: 5’-GAAGAGAGGCATGTTGGAGACTTGCGCAATGTGACCGCGGACAAAGATG-3’ G93S: 5’-

GCAATGTGACCGCGGACAAAGATTCTGTGGCCGATGTATCGATTGAAGAT -3’ and the reverse primer 5’-CGATCCCAATTACACCACAAGCCA -3’. In the case of the L144F mutation, the following respective forward and reverse primers were used: 5’-GACCAGTGAAGGTGTGGGGAAG-3’ and 5’-GGCGATCCCAATTACACCACAAGCGAAACGGGATCCAGCGTTTCCTGTC -3’. Alanine was added at the C-terminus using the respective and reverse forward primers: 5’-AAGGAGATATACATATGAATAAGGCAAAAACTTTACTCTTCAC -3’ and 5’-TTGCTGGTTTACCGGTTTACGCTTGGGCGATCCCAATTACACCA-3’. The wild-type SOD1gene was cloned into the pRFS-Duet plasmid using the respective forward and reverse primers:

5’-AATTTTGTTTAACTTTAATAAGGAGATATACCATGAATAAGGCAAAAACTTTAC TCT-3’ and 5’-ACTTAAGCATTATGCGGCCGCAAGCTTTTATTGGGCGATCCCAATTACAC-3’. The SOD1genes with disease-causing mutations were cloned into the pRFS-Duet plasmid using the respective forward and reverse primers:

5’-AAGGAGATATACATATGAATAAGGCAAAAACTTTACTCTTCAC -3’ and 5’-TTGCTGGTTTACCGGTTTATTGGGCGATCCCAATTACAC-3’.

### Sample preparation

A single *E. coli* BL21 (DE3) colony harboring the pRSF-Duet plasmid containing the wild-type and mutant SOD1 genes was picked and used to inoculate 5 mL LB medium containing 50 µg ml^-1^ kanamycin. The cells were grown for ∼8 hours at 37 °C and 0.4 ml of the culture were then transferred to 50 mL 2x YT medium containing 50 µg ml^-1^ kanamycin and growth was continued at 37 °C until an OD_600nm_ of ∼0.7 was reached. Protein expression was then induced by adding 0.25 mM IPTG, 0.5 mM CuSO_4_ and 0.01 mM ZnCl_2_ to the medium and growth was continued for 12 hours at 25 °C. Harvesting was carried out by centrifuging 50 ml of the cell cultures at 4,000 g for 15 min. Cell pellets were washed once with water to remove residual growth medium and then stored at -80 °C until use. Cells were then resuspended in 2 ml of 1 M ammonium acetate buffer (pH 7.0) containing protease inhibitors (0.3 mM PMSF, 1 mM benzamidine and 1.1 µg ml^-1^ pepstatin A) and lysed by sonication. The lysate was cleared by centrifugation at 11,000 g for 15 min and the supernatant was used for MS analyses.

### Native mass spectrometry analysis

Measurements were performed using a Q Exactive Plus Orbitrap mass spectrometer (Thermo Fisher Scientific, Bremen, Germany) modified for optimal transmission and detection of high molecular weight ions (34). The samples were diluted 100-fold with 1 M ammonium acetate buffer (pH 7.0) containing 40 mM imidazole before MS analysis. Typically, a sample of 2 µl was loaded into a gold-coated nano-ESI capillary prepared in-house as previously described (35) and then sprayed into the instrument. Conditions within the mass spectrometer were adjusted to preserve non-covalent interactions with the source operating in positive mode. The following experimental parameters were used: capillary voltage, 1.7 kV; HCD direct voltage, 35 eV; and an inlet capillary temperature of 180 °C. MS spectra were recorded at a resolution of 17,500 (at m/z 200). The areas of the peaks in each mass spectrum, which correspond to the concentrations of the different complexes (taking into consideration the adherence of adducts), were calculated using Kaleidagraph version 4.5 and OriginPro 8. The analysis was performed for charge state 11^+^. For each double-mutant cycle, three biological replicates were prepared and analyzed and at least three independent measurements were acquired for each replica.

## Supporting information

Supplemental Figues 1 and 2

## Acknowledgements

Part of this work was carried out while E.M.M. was an Erna and Jakob Michael Visiting Professor at the Weizmann Institute. A.H. is grateful for support from the Helen & Milton A. Kimmelman Center for Biomolecular Structure and Assembly and the Ilse Katz Institute for Material Sciences and Magnetic Resonance Research. A.H. is an incumbent of the Carl and Dorothy Bennett Professorial Chair in Biochemistry. M.S. is grateful for the financial support of a starting grant from the European Research Council (Horizon 2020) no. 636752 and an Israel Science Foundation grant no. 300/17. M.S. is the incumbent of the Aharon and Ephraim Katzir Memorial Professorial Chair.

## Notes

### Competing Interest Statement

The authors have declared no competing interest.

## References

1. Levy ED, Teichmann S. 2013. Structural, evolutionary and assembly principles of protein oligomerization. Prog. Mol. Biol. Transl. Sci. 117, 25–51.

2. Bergendahl LT, Gerasimavicius L, Miles J, Macdonald L, Wells JN, Welburn JPI, Marsh JA. 2019. The role of protein complexes in human genetic disease. Protein Sci. 28, 1400–1411.

3. Klein C, Westenberger A. 2012. Genetics of Parkinson’s disease. Cold Spring Harb. Perspect. Med. 2, a008888.

4. Beaver SK, Mesa-Torres N, Pey AL, Timson DJ. 2019. NQO1: A target for the treatment of cancer and neurological diseases, and a model to understand loss of function disease mechanisms. Biochim. Biophys. Acta 1867, 663–676.

5. Taylor JP, Brown RH Jr, Cleveland DW. 2016. Decoding ALS: from genes to mechanism. Nature 539, 197–206.

6. Rosen DR, Siddique T, Patterson D, Figlewicz DA, Sapp P, Hentati A, Donaldson D, Goto J, O’Regan JP, Deng HX, et al. 1993. Mutations in Cu/Zn superoxide dismutase gene are associated with familial amyotrophic lateral sclerosis. Nature 362, 59–62.

7. Wang Q, Johnson JL, Agar NY, Agar JN. 2008. Protein aggregation and protein instability govern familial amyotrophic lateral sclerosis patient survival. PLoS Biol. 6, e170.

8. Horovitz A, Fleisher RC, Mondal T. 2019. Double-mutant cycles: new directions and applications. Curr. Opin. Struct. Biol. 58, 10–17.

9. Sokolovski M, Cveticanin J, Hayoun D, Korobko I, Sharon M, Horovitz A. 2017. Measuring inter-protein pairwise interaction energies from a single native mass spectrum by double-mutant cycle analysis. Nat. Commun. 8, 212.

10. Lössl P, van de Waterbeemd M, Heck AJ. 2016. The diverse and expanding role of mass spectrometry in structural and molecular biology. EMBO J. 35, 2634–2657.

11. Liko I, Allison TM, Hopper JT, Robinson CV. 2016. Mass spectrometry guided structural biology. Curr. Opin. Struct. Biol. 40, 136–144.

12. Chandler SA, Benesch JL. 2018. Mass spectrometry beyond the native state. Curr. Opin. Chem. Biol. 42, 130–137.

13. Cveticanin J, Netzer R, Arkind G, Fleishman SJ, Horovitz A, Sharon M. 2018. Estimating interprotein pairwise interaction energies in cell lysates from a single native mass spectrum. Anal. Chem. 90, 10090–10094.

14. Wright GSA, Antonyuk SV, Hasnain SS. 2019. The biophysics of superoxide dismutase-1 and amytrophic lateral sclerosis. Q. Rev. Biophys. 52, e12.

15. Nordlund A, Oliveberg M. 2008. SOD1-associated ALS: a promising system for elucidating the origin of protein-misfolding disease. HFSP J. 2, 354–364.

16. Redler RL, Dokholyan NV. 2012. The complex molecular biology of amyotrophic lateral sclerosis. Prog. Mol. Biol. Transl. Sci. 107, 215–262.

17. Shi Y, Acerson MJ, Abdolvahabi A, Mowery RA, Shaw BF. 2016. Gibbs energy of superoxide dismutase heterodimerization accounts for variable survival in amyotrophic lateral sclerosis. J. Am. Chem. Soc. 138, 5351–5562.

18. Lindberg MJ, Byström R, Boknäs N, Andersen PM, Oliveberg M. 2005. Systematically perturbed folding patterns of amyotrophic lateral sclerosis (ALS)-associated SOD1 mutants. Proc. Natl. Acad. Sci. USA 102, 9754–9759.

19. Borchelt DR, Lee MK, Slunt HS, Guarnieri M, Xu ZS, Wong PC, Brown RH Jr, Price DL, Sisodia SS, Cleveland DW. 1994. Superoxide dismutase 1 with mutations linked to familial amyotrophic lateral sclerosis possesses significant activity. Proc. Natl. Acad. Sci. USA 91, 8292–8296.

20. Witan H, Kern A, Koziollek-Drechsler I, Wade R, Behl C, Clement AM. 2008. Heterodimer formation of wild-type and amyotrophic lateral sclerosis-causing mutant Cu/Zn-superoxide dismutase induces toxicity independent of protein aggregation. Hum. Mol. Genet. 17, 1373–1385.

21. Brasil AA, de Carvalho MDC, Gerhardt E, Queiroz DD, Pereira MD, Outeiro TF, Eleutherio ECA. 2019. Characterization of the activity, aggregation, and toxicity of heterodimers of WT and ALS-associated mutant Sod1. Proc. Natl. Acad. Sci. USA 116, 25991–26000.

22. Xu G, Ayers JI, Roberts BL, Brown H, Fromholt S, Green C, Borchelt DR. 2015. Direct and indirect mechanisms for wild-type SOD1 to enhance the toxicity of mutant SOD1 in bigenic transgenic mice. Hum. Mol. Genet. 24, 1019–10135.

23. Prudencio M, Durazo A, Whitelegge JP, Borchelt DR. 2010. An examination of wild-type SOD1 in modulating the toxicity and aggregation of ALS-associated mutant SOD1. Hum. Mol. Genet. 19, 4774–4789.

24. Rumfeldt JA, Stathopulos PB, Chakrabarrty A, Lepock JR, Meiering EM. 2006. Mechanism and thermodynamics of guanidinium chloride-induced denaturation of ALS-associated mutant Cu,Zn superoxide dismutases. J. Mol. Biol. 355, 106–123.

25. Elam JS, Taylor AB, Strange R, Antonyuk S, Doucette PA, Rodriguez JA, Hasnain SS, Hayward LJ, Valentine JS, Yeates TO, Hart PJ. 2003. Amyloid-like filaments and water-filled nanotubes formed by SOD1 mutant proteins linked to familial ALS. Nat. Struct. Biol. 10, 461–467.

26. Banci L, Bertini I, D’Amelio N, Libralesso E, Turano P, Valentine JS. 2007. Metalation of the amyotrophic lateral sclerosis mutant glycine 37 to arginine superoxide dismutase (SOD1) apoprotein restores its structural and dynamical properties in solution to those of metalated wild-type SOD1. Biochemistry 46, 9953–9962.

27. Broom HR, Rumfeldt JA, Vassall KA, Meiering EM. 2015 Destabilization of the dimer interface is a common consequence of diverse ALS-associated mutations in metal free SOD1. Protein Sci. 24, 2081–2089.

28. Khare SD, Caplow M, Dokholyan NV. 2006. FALS mutations in Cu, Zn superoxide dismutase destabilize the dimer and increase dimer dissociation propensity: a large scale thermodynamic analysis. Amyloid 13, 226–235.

29. Kato S, Shimoda M, Watanabe Y, Nakashima K, Takahashi K, Ohama E. 1996. Familial amyotrophic lateral sclerosis with a two base pair deletion in superoxide dismutase 1: gene multisystem degeneration with intracytoplasmic hyaline inclusions in astrocytes. J. Neuropathol. Exp. Neurol. 55, 1089–1101.

30. Yamagata K, Furuta H, Oda N, Kaisaki PJ, Menzel S, Cox NJ, Fajans SS, Signorini S, Stoffel M, Bell GI. 1996. Mutations in the hepatocyte nuclear factor-4? gene in maturity-onset diabetes of the young (MODY1). Nature 384, 458–460.

31. Unger T, Jacobovitch Y, Dantes A, Bernheim, R, Peleg Y. 2010. Applications of the Restriction Free (RF) cloning procedure for molecular manipulations and protein expression. J. Struct. Biol. 172, 34–44.

32. Getzoff ED, Cabelli DE, Fisher CL, Parge HE, Viezzoli MS, Banci L, Hallewell RA. 1992. Faster superoxide dismutase mutants designed by enhancing electrostatic guidance. Nature 358, 347–351.

33. Sekhar A, Rumfeldt JAO, Broom HR, Doyle CM, Sobering RE, Meiering EM, Kay LE. 2016. Probing the free energy landscapes of ALS disease mutants of SOD1 by NMR spectroscopy. Proc. Natl. Acad. Sci. USA 113, E6939–E6945.

34. Ben-Nissan G, Belov ME, Morgenstern D, Levin Y, Dym O, Arkind G, Lipson C, Makarov AA, Sharon M. 2017. Triple-stage mass spectrometry unravels the heterogeneity of an endogenous protein complex. Anal. Chem. 89, 4708–4715.

35. Kirshenbaum N, Michaelevski I, Sharon, M. 2010. Analyzing large protein complexes by structural mass spectrometry. J. Vis. Exp., e1954.

