## Supplemental Figues 1 and 2 for "Insight into the autosomal-dominant inheritance pattern of SOD1-associated ALS from native mass spectrometry"

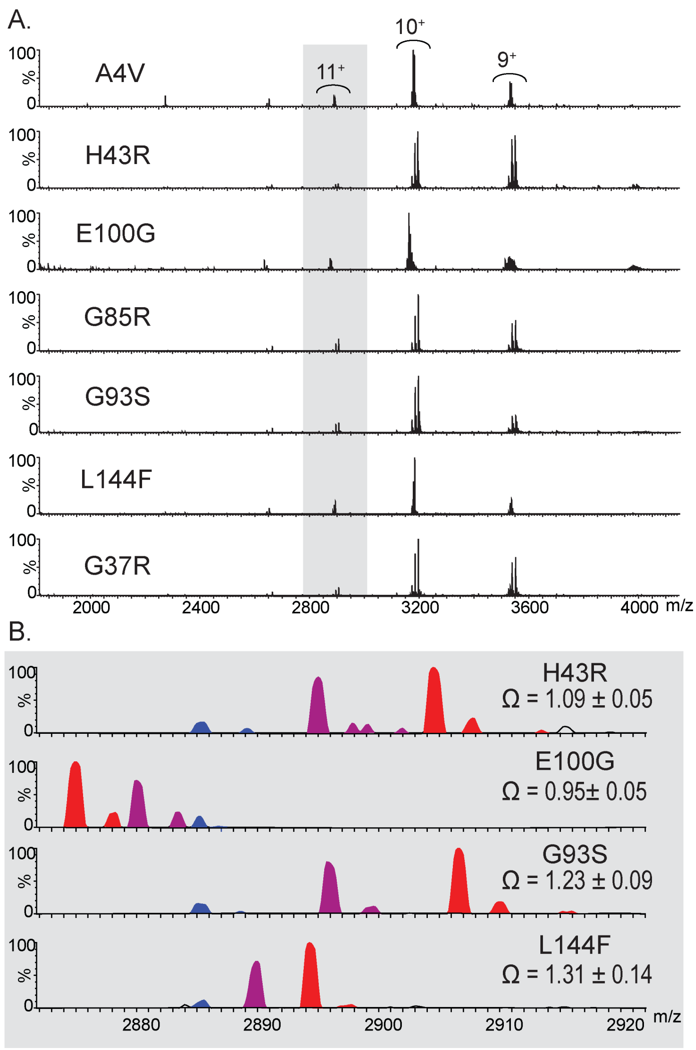


**Figure S1**. Native MS measurements of the wild-type and mutant SOD1 homo- and heterodimers in crude cell lysates. (A) The mass spectra of lysates containing co-expressed wild-type SOD1 and the different mutants are shown with the peaks for $11^+$ charge state highlighted in gray. (B) The mass spectra of the $11^+$ charge state are shown for lysates containing co-expressed wild-type SOD1 and either the H43R, E100G, G93S or L114F mutants. The peaks for the wt/wt, mut/mut and mut/wt (and wt/mut) dimers are shown in blue, red and purple, respectively.

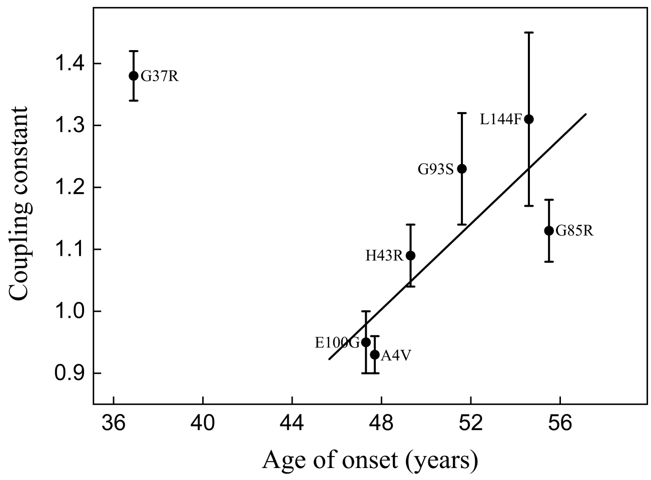


**Figure S2**. Plot of the coupling energies between wild-type and mutant SOD subunits and the ages of onset associated with the respective mutation. For simplicity, the data (excluding that for G37R) were subjected to a linear fit ($r^2$ = 0.64; P = 0.056) although the underlying relationship may be monotonically increasing but nonlinear. The ages of onset were taken from ref. 7.
